# Research on the temporal evolution track and influence of green development in the last decade

**DOI:** 10.1101/2022.01.20.477152

**Authors:** Yunfang Chen, Qingkui Lai, Xuan Ji, Cunyu zhang, Mengxiong Xiao

## Abstract

Green development is important to realize the coordinated development of local ecological environment, economy and society, but also an important way to promote the harmony between human and land. This study utilizes projection pursuit model (PPM) measure, as well as the system gray prediction model GM (1, N), and other measurement methods, build an evaluation system from three dimensions of green fortune, green growth and green benefits to analyze the temporal evolution, development trend and its influencing factors of Baoshan’s ecological engineering of project acres, the results show: (1)The overall level of the green development system in Baoshan, Yunnan Province, is relatively good and shows an increasing trend year by year, with an average growth rate of 18.3%, and there is a synergistic coupling effect among the three subsystems. (2) Regional green development has achieved remarkable progress, but problems and pressure among the three subsystems are still grim. The sub-system of green growth and green benefits declined from 2010 to 2016, with an annual decline rate of 12%, and then picked up slightly in the next three years, reflecting the pressure of resources and ecological environment in this region and the difficulty of the sub-system of green wealth growth. (3) The prediction analysis shows that the green development index in Baoshan, Yunnan Province will keep a steady rise in the next two years, and the predicted value will be 4.51 in 2020, with an average growth rate of 15.1%, and the regional green development system is in good condition. (4) Green development is mainly affected by environmental pollution, industrial structure, urbanization, population, market and other factors from the three subsystems of local ecology, ecology, economy and social benefits. This paper puts forward suggestions in terms of popularizing the concept of green development, improving the level of ecological economic development and promoting the regional environment bearing capacity, green development model of ecological engineering area is also built. It aims to have some practical reference value for prompting regional ecological civilization construction and green development. And provide regional support for the post-2020 Global Biodiversity Framework and the Achievement of the United Nations Sustainable Development Goals.

## Introduction

China confronts with severe resource and environment pressure now, in addition, special population, resource and environmental condition decide China must implement the strategy of sustainable development. The White Paper on Chinese Population, Environment and Development in the 21st Century is the first sustainable development strategy in China. Then national 10th Five-Year plan, 11th Five-year plan promote the implement of the strategy constantly and they call for building a resource-conserving and environment friendly society. It is the 12th Five-Year Plan that green development is put forwards and it advocates “establishing the concept of green and low-carbon development”. The 13th Five-Year-Plan put forward that green is a necessary condition for sustainable development and important embodiment of people’s pursuit of a better life. The report to 19th National Congress of the Communist Party of China (CPC) pointed that we should accelerate the reform of the ecological civilization system to better meet growing ecological needs of the people; it also emphasizes the strategic development direction of realizing excellent ecological environment through green development. Up to now, China’s green development has gradually changed from simple to complex, from one-dimension to multi-dimension exploration.

Scholars at home and abroad have had many theoretical explorations on the concept and connotation, dynamic and static assessment, driving forces and realization approaches of green development have also been explored, but there is major issue to be addressed. Firstly, the concept of green development in China is diversified and limited due to the combination of theory and practice from multiple disciplines and perspectives; secondly, it is believed that green development is ecological protection, which is confused with the coordinated development of economy and society, the evaluation of economic and social development; which leads to the fuzzy evaluation of green development; to understand green development from a perspective of green economy, industrial greening or ecological system, resource system and energy system, which lacks a systematic and overall understanding of China’s green development. It is essential to clarify the connotation and theoretical system of green development and construct an evaluation system for regional green development. From the above mentioned, the world has come into a new era with the theme of “green development”, green development has gradually become the inevitable choice for many countries and regions to confront the financial crisis, environment and resource challenges and sustainable development. Based on the scientific understanding of green development system and connotation, this paper constructs an evaluation system for regional green development accordingly.

In figure 1, located in the southwest of Yunnan, to the west is Myanmar, adjoin Dehong, Baoshan is an essential route since ancient times. From 2016, in accordance with the development direction of “Three Green Card” of Yunnan Government, Baoshan has constructed the ecological engineering project in combination with its regional characteristics and urban ecological development strategy to integrate the elements of “mountain, water and city” into one, and draw a new blueprint for regional ecological development. The implementation of ecological engineering project will promote the development of ecological civilization in Yunnan, but also improve the living standards of local residents and radiates the economic development of surrounding areas. So this paper studies the change of the green development index and green development level, evolution mechanism under the promotion of ecological engineering project in recent 10 years in Longyang District, Baoshan, to provide reference for the promotion of green development and ecological civilization construction in Yunnan Province.

**Figure1.**
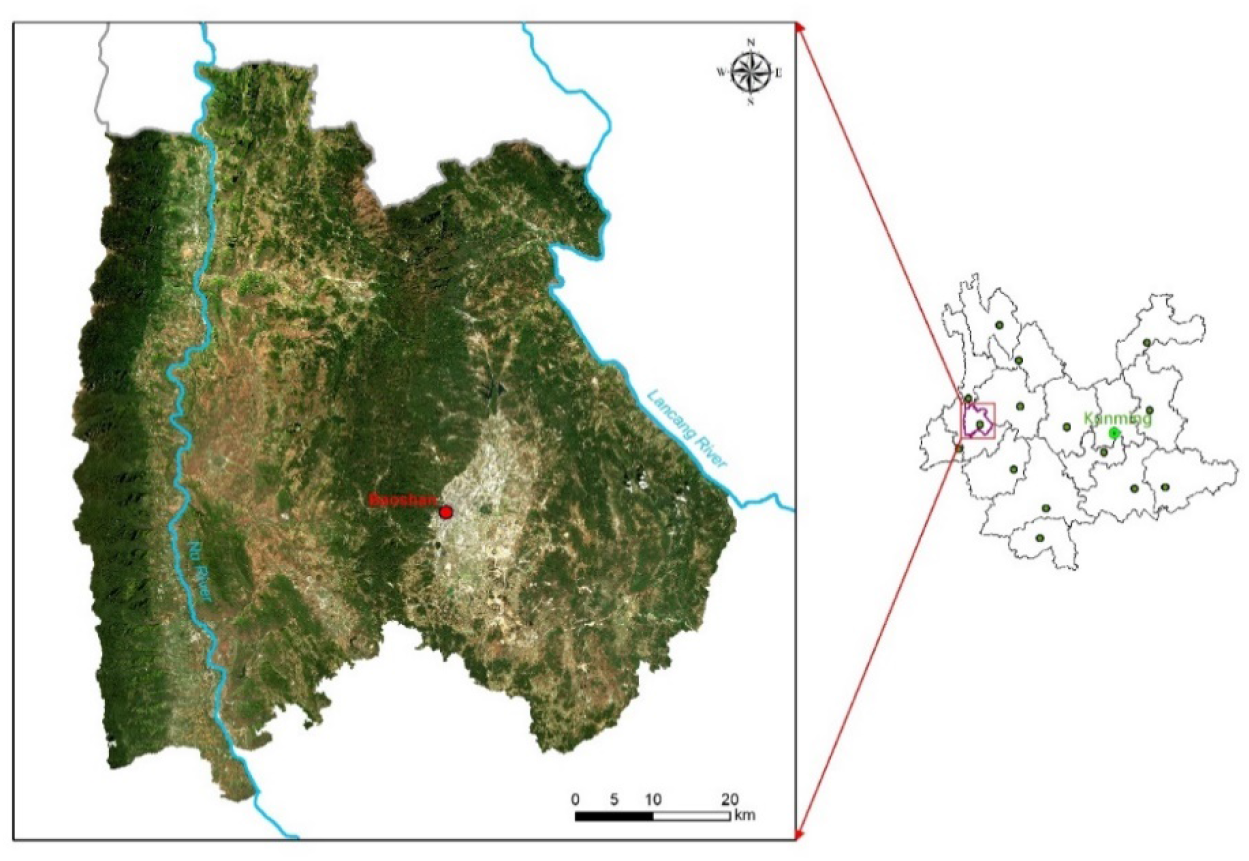
Sketch map of study area

### The measurement method of Green Development Index

#### The basic connotation and main content of Green Development

The essence of green development realizing the concept of sustainable development and widely recognized by the international community (Liu Mingguang, 2017). The relationship between them is the same strain, it is not only the inheritance of sustainable development, but also the theoretical innovation of sustainable development in China. It is also a significant theoretical contribution to China’s response to the objective reality of global environment deterioration, which is not a single protection and construction of the ecological environment, but an organic combination of three subsystems of natural ecology, economy and society, green development is a development path with high efficiency and coordination of ecology, economy and has systematic integrity, dynamism and ecology and society, which aims to increase human welfare and accumulate green wealth, the economic behavior of low consumption, low emission and reasonable consumption, and the continuous enrichment of ecological capital as the main features, and the purpose is to achieve the harmonious coexistence and mutual benefit between human and nature. Green development is the requirement for sustainable development, an important manifestation of Chinese people’s pursuit of a better life, and a fundamental approach to Chinese ecological civilization construction.

The connotation of green development varies from one to another, but it is basically the same connotation in the academic circle: that is focusing on the process of regional high-quality development, namely development efficiency and innovation-driven (Lorek, 2014). Maximize ecological, economic and social benefits in the process of high-quality development. (Loiseau, 2016; Amato, 2017), thus, the optimal development mode of coordinated regional resource and environment system, economic system and social system is an important path to realize green development.

Respect for ecological system, economic system and social system is the requirement for green development, which covers 3 areas: first, to realize the green restructuring of resource elements allocation, promote the transition of green production function; second, to realize the decoupling of economic growth and economic development speed from ecological deficit; third, establish the development concept of “unity of nature and man” and mutual benefit of nature and man (Hu Angang, 2015). From the perspective of system and based on system function, mechanism, development strategy of green development were analyzed, and the concept of “Three Circles” model is set up, the green development is regarded as the second generation of sustainable development including green wealth, green growth and green benefits. Among them, green wealth is a foundation, green growth is means and green welfare is the target (Hu Angang, 2014). therefore, green development is a kind of relief efficiency mode of sustainable development, in coordination with economic and social development of efficient environment symbiotic for development direction, by realizing the transformation of industrial environment friendly resource saving within the regional system, improving the cleaner production mechanism and changing the production process (Chu Dajian, 2012), with the ultimate purpose for achieving green development and high-level development of ecological civilization in the region.

#### The evaluation system of Green Development

In recent years, with the extensive application of green economy and sustainable development, the evaluation index system has been continuously deepened, mainly including environmental sustainability, inclusiveness, green economy, etc.

The research on the evaluation of green development mainly covers two aspects: evaluation and construction of green development measurement index system. The methods of green development evaluation include input-output and cost-benefit model, entropy method (Guo Fuyou et al, 2021), spatial autocorrelation analysis (Cheng Yu et al, 2019), LMDI measurement model (Carfi, 2012), environmental carrying capacity, etc. In terms of indicator selection and system construction of green development, most of the indicator systems are selected by referring to environment carrying capacity index, green industry, green economy evaluation, etc., or the green GDP of resource and environmental cost and ecological efficiency index are calculated to measure the efficiency of green development (Li Lin, 2012; Zhao Liang et al, 2016). For example, the green development index was established based on the human development index, and the green development index of 123 countries and regions was measured (Li Xiaoxi et al., 2014). Some also analyzed the change of the time pattern of green economy by calculating China’s green CDP (Shen Xiaoyan et al., 2017). Scholars studied the spatial evaluation model of China’s green development efficiency from the perspective of resource and environment by using the method of data envelopment analysis (Qian Zhengming and Liu Xiaochen, 2014).

As for green development index, Yang Duogui (2006) built evaluation system by analyzing environment benefits, energy consumption, the metabolism of environment, environment pollution, resource consumption. Green economy development index is also established according to the principle of ecosystem material flow, containing three aspects: green production, green consumption and green health (Xiang Shujian, 2012); besides, based on green production, living, green environment and green new policy, studies have constructed the evaluation index system of green development at the provincial level(Liu Mingguang, 2017).

The previous research on green development in China focused on the evaluation of green economy, industrial greening degree and sustainable development degree that is necessary to conduct a systematic study on the current situation and deficiency of green development, and conduct a systematic evaluation, condition simulation, and development trajectory analysis of green development, which is of great practical significance to the regional green development and transformation.

Based on the green development concept put forward by Hu Angang, this study constructs the evaluation index system of green development according to its green growth, green wealth and green welfare. Referenced the current research results and regional green development strategy design, the regional green development evaluation index system is designed according to the scientific, regional, operable and systematic comprehensive principles (Wang Yong, Li Haiying, 2018). This study analyze green development evaluation system including three layers: the first layer for the total system, namely the development of regional green evaluation index system; the second is subsystem layer, or layer evaluation of ecological economic and social subsystem; the third layer of elements, including green wealth, green growth and green welfare and the fourth layer of index layer, with the total of 25 indicators. Among them, green wealth mainly includes: 7 basic indicators such as annual afforestation area, green coverage rate, per capital arable land, regional emission of SO^2-^; the green increase covers 8 basic indicators such as per capita GDP, energy consumption and land productivity; green welfare includes 10 basic indicators such as the numbers of students in this area, the area’s average income, per capita public green land, and effective irrigation area (Table 1 below).

**Table1.**
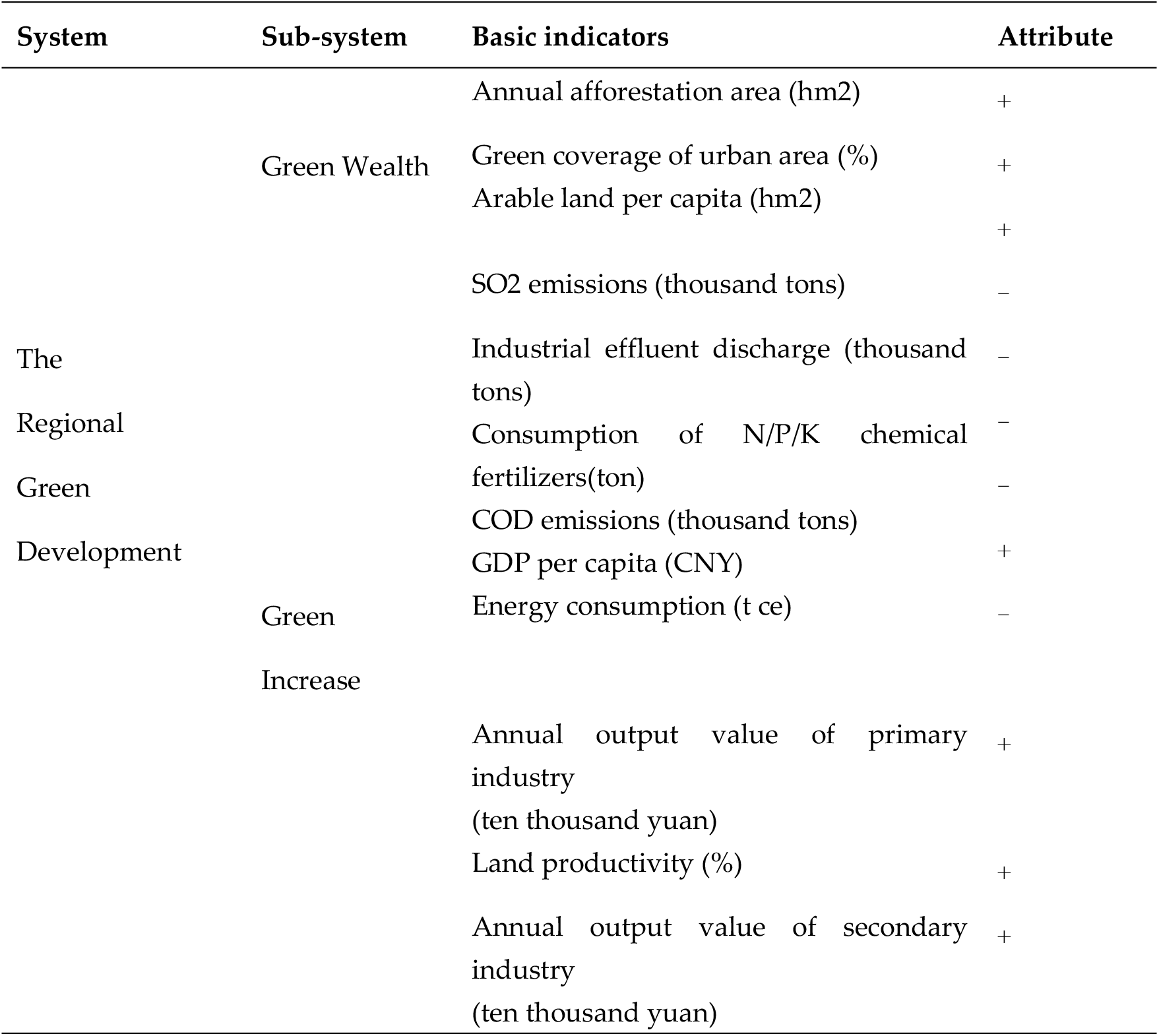
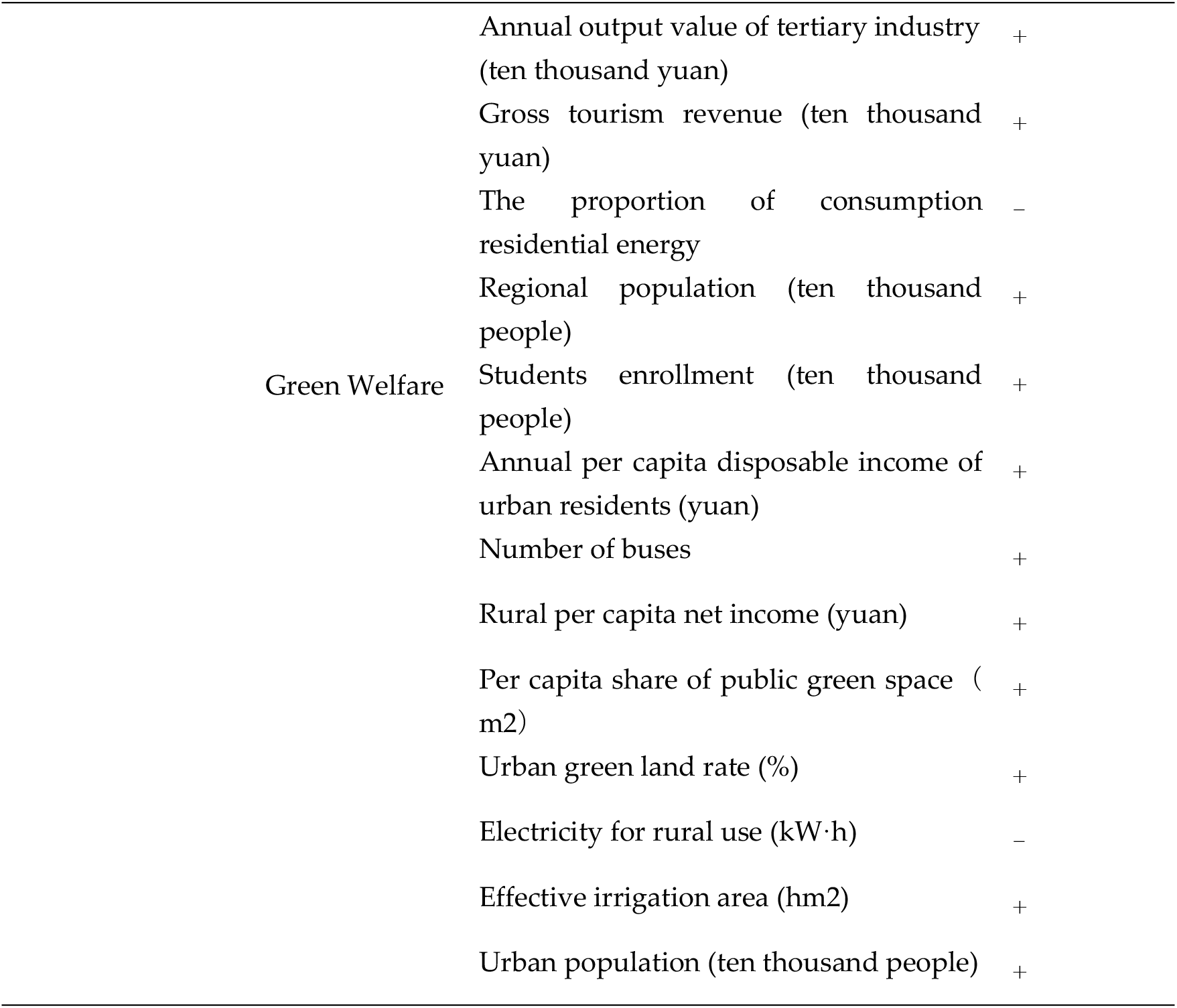
The construction of regional green-development

#### Method of Regional Green development Index Measurement

The projection pursuit model (PPM) is used to evaluate the regional green development, which can process the nonlinear and multi-dimensional data scientifically and systematically, and can effectively calculate the regional green development, the multivariate complex system evaluation problem. By using the projection optimization function to reduce the dimension of the multi-dimension data, and use the low-dimensional data to simulate and analyze the characteristics of the multi-dimensional data and the projection value, then the synthesis problem of the multi-dimensional data.

① Data standardization

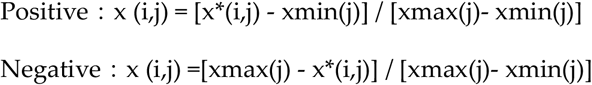
② Projection function Suppose n={n(1), n(2), …, n(q)} is the projection direction vector, and the one-dimensional projection value of region i :

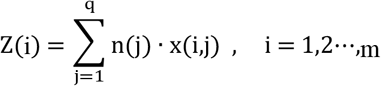 Then project objective function:

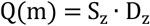 In the above equation, Sz is the standard deviation of Z(i); Dz is the local density of Z (i).

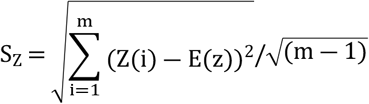
③ Optimization of Projection Objective Function

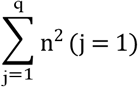 The optimization gist of the projection is a nonlinear optimization problem with the variable: {n(j)/j=1, 2, …, q}, and the maximum value can be processed by using accelerate genetic algorithm. Parameters as follows: initial 400, crossover probability 0.8, accelerate times:50.
④ The optimal value. The optimal projection direction (i.e., weight) n* is multiplied by the normalized index value, and the projection value X is(i) obtained by summing, namely, the evaluation score of green development index is obtained. The larger the projection is, the higher the score is:

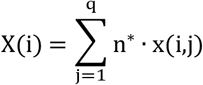

#### The main source of indicator data

Based on the above evaluation index system of regional green development, the research takes Baoshan, Yunnan, a project area of 10.000 mu as the research object, and the basic index derives from the Statistical Yearbook of Baoshan City, Yunnan, Statistical Yearbook of Longyang District, Baoshan City, Yunnan, and the environmental Statistical data of the study area from 2010-2019.

### The green development and evolution track of ecological engineering construction area

#### Temporal evolution track analysis of green development

The projection pursuit model (PPM) was used to measure the green development validity of 10.000 mu project of Baoshan in Yunnan province. The results are shown in Table 2 below. From the perspective of the overall development system of this region, the overall green development level shows the trend of increasing. As shown in Figure 2, from 1.31 in 2010 to 3.71 in 2019, the average annual growth rate is about 18.3%, indicating that the level of green development in this region has been significantly improved. Realizing the great effect of resource environment on the regional economic and social development, local government must follow the natural ecology of the regional development pattern and economic laws, make the green development as a higher development strategy, focus on the coordinated development of local ecological environment, economy and society to optimize the sustainable development of regional human and land system. The government of Baoshan make great effort to build an ecological corridor project covering an area of more than 10.000 mu, actively promote the strategies of building new urbanization, rural revitalization of the demonstration zone, beautiful and livable city of Baoshan, promoting regional across development and speeding up the pace of urban ecological development and the sustainable development of local production, consumption structure and the system structure optimized constantly, strengthen urban environmental governance, reduce resource consumption and improve the level of ecological progress. However, the evaluation of local green development system shows that the overall growth rate of green development in China is growing slowly, and the concept of sustainable development needs to be internalized into the comprehensive decision-making of local top-level design, planning and development.

**Table2.**
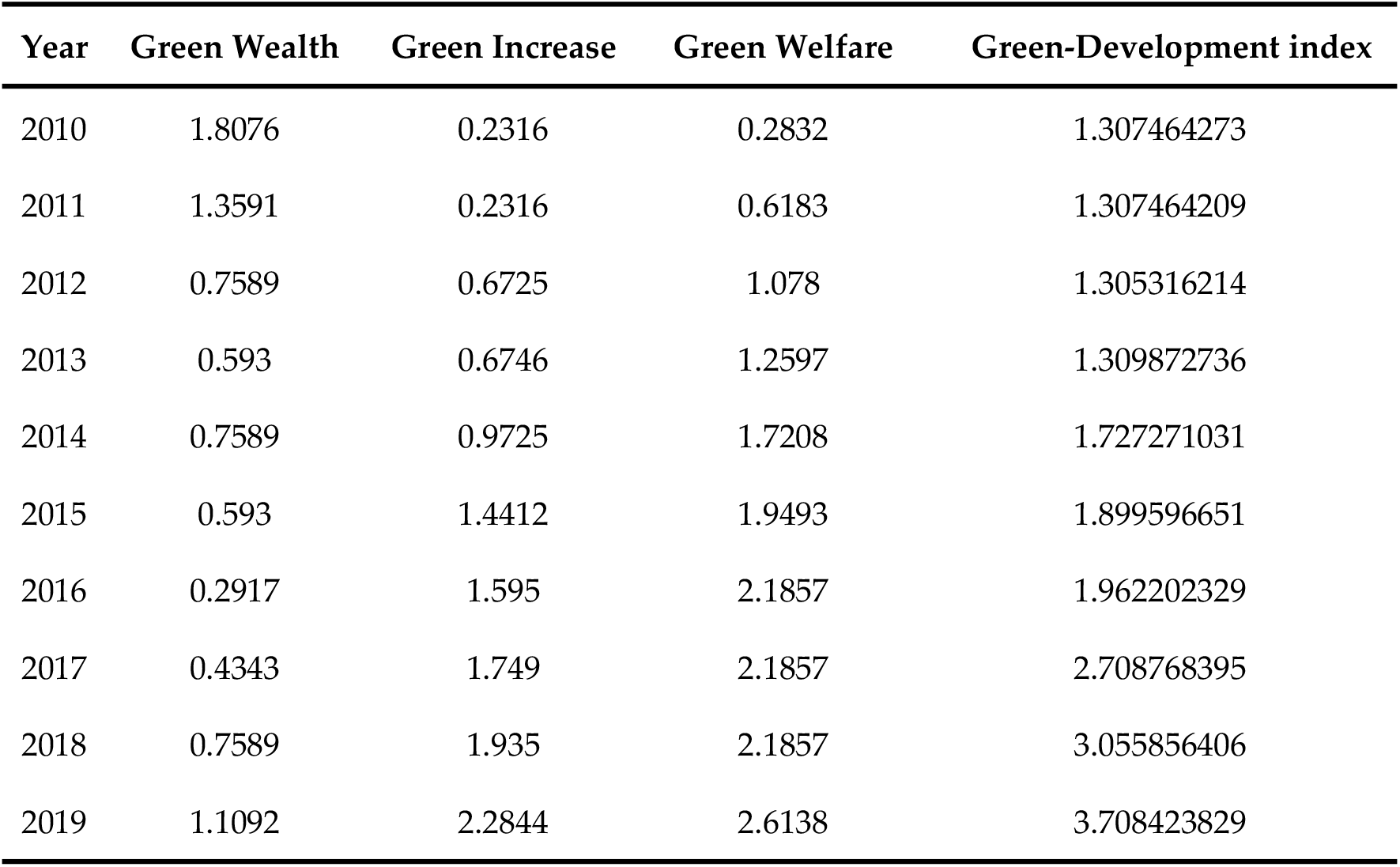
Study area development index and its composition from 2010-2019

| Year | Green Wealth | Green Increase | Green Welfare | Green-Development index |
| --- | --- | --- | --- | --- |
| 2010 | 1.8076 | 0.2316 | 0.2832 | 1.307464273 |
| 2011 | 1.3591 | 0.2316 | 0.6183 | 1.307464209 |
| 2012 | 0.7589 | 0.6725 | 1.078 | 1.305316214 |
| 2013 | 0.593 | 0.6746 | 1.2597 | 1.309872736 |
| 2014 | 0.7589 | 0.9725 | 1.7208 | 1.727271031 |
| 2015 | 0.593 | 1.4412 | 1.9493 | 1.899596651 |
| 2016 | 0.2917 | 1.595 | 2.1857 | 1.962202329 |
| 2017 | 0.4343 | 1.749 | 2.1857 | 2.708768395 |
| 2018 | 0.7589 | 1.935 | 2.1857 | 3.055856406 |
| 2019 | 1.1092 | 2.2844 | 2.6138 | 3.708423829 |

**Figure2.**
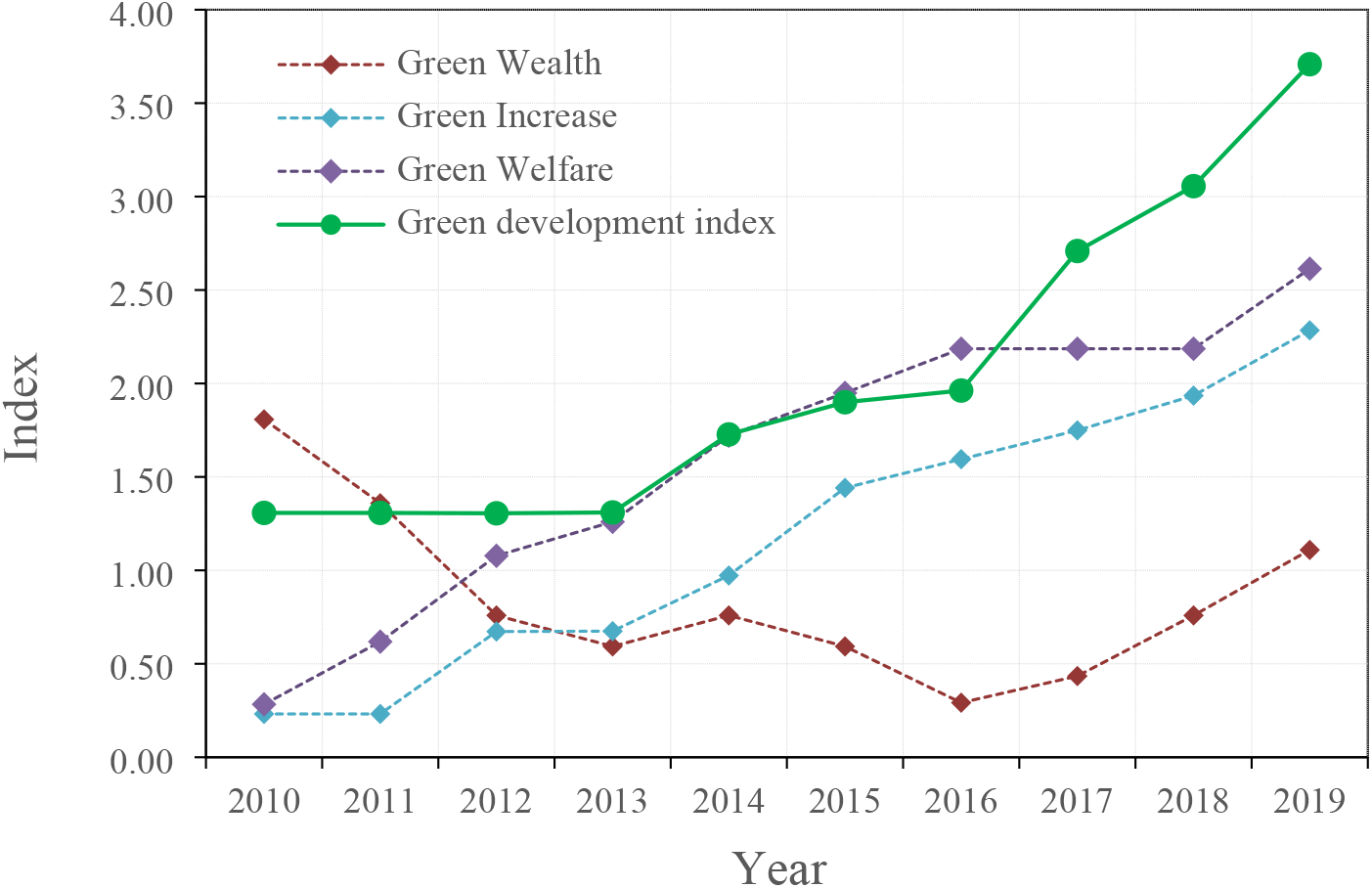
Green-development index of Baoshan’s ecological engineering of Project Area

**Figure2.**
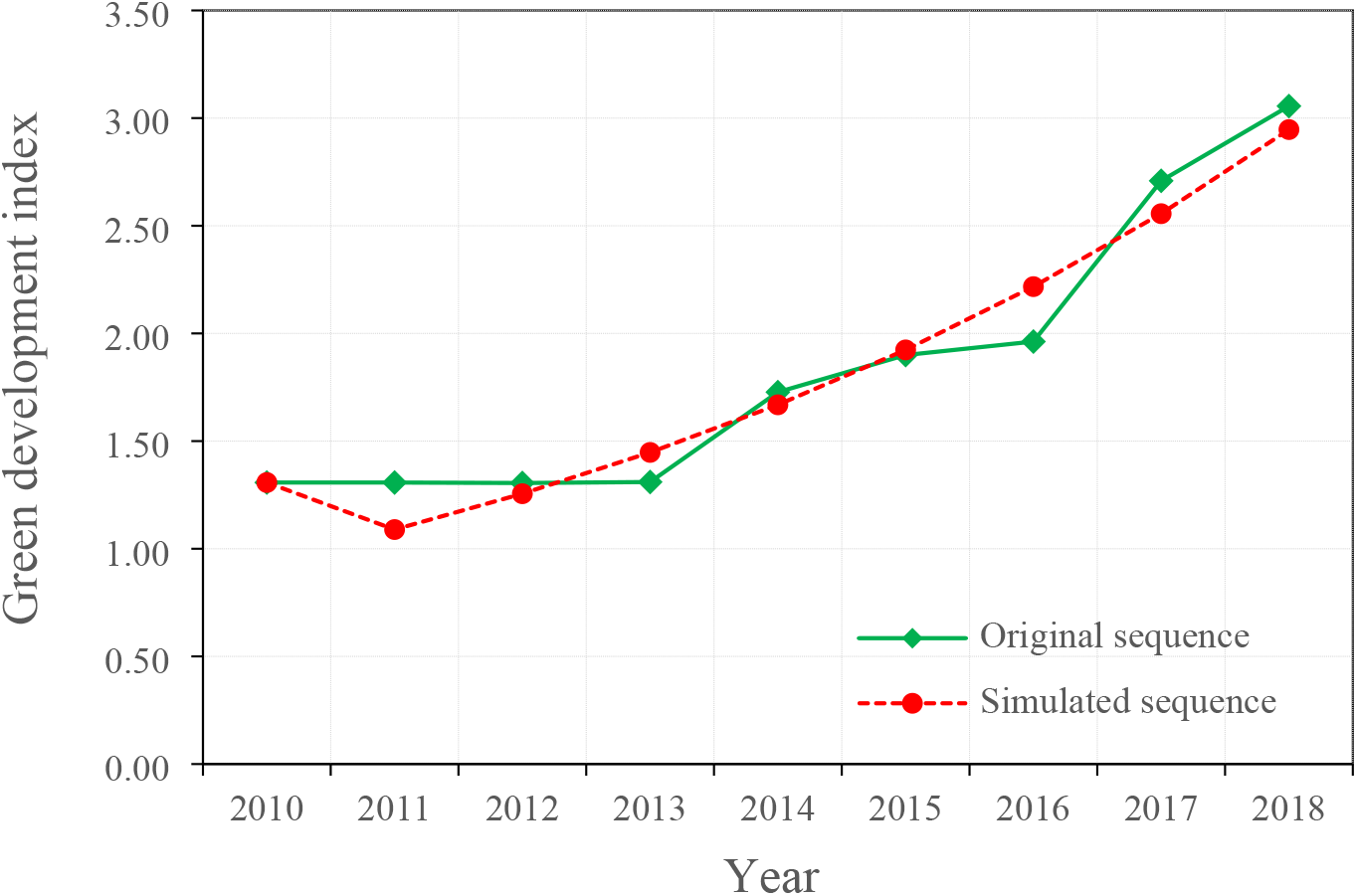
Dynamic simulation analysis of Green Development of Baoshan’s ecological engineering Area from 2010 to 2018

From the point of the structure of the green development assessment system in the construction area, the “green wealth” subsystem was in a declining state at this stage. From 2010 to 2016, the “green wealth” index, which represents the regional ecosystem and environmental pressure, continued to decline from 1.81 in 2010 to 0.29 in 2016, with an annual decline rate of 12%. Therefore, restore wetlands and strengthen ecological management, Baoshan in Yunnan province has implemented the “Three Ten thousand mu Project” since 2016, that is, the ten thousand mu wetland restoration project, the ten thousand mu sightseeing agriculture restoration project and the ten thousand mu ecological management vegetation restoration project in Dongshan, integrating the elements of mountains, water, fields, parks and towns to promote the ecological development of regional cities. The development index of the green wealth subsystem in the construction area of 10,000 mu has been relatively significantly recovered from 2017 to 2019, rising from 0.29 in 2016 to 1.11 in 2019. It shows that during the study period, local governments actively respond to national policies, strengthen environmental governance investment and management, promote green industrial transformation, and promote ecological red line strategy, so that the uncontrolled development and utilization of regional resources and environment and the disorderly expansion of land space are gradually under effective control. The discharge of SO^2^, industrial wastewater, the application amount of nitrogen, phosphorus, potassium and compound fertilizer, and the discharge of COD in the study area have been continuously reduced, and the ecological and environmental functions of the region have been significantly restored, indicating that the resource consumption intensity, pollutant discharge and economic scale growth of the region gradually decoupled, but slightly slow.

With the continuous improvement of the regional economic development level, economic subsystem, the involvement of the structure in the direction of rationalization, ordering degree of green industry economy also gradually improve, the subsystem of “green growth” representing economic development and the subsystem of “green welfare” representing social development elements both show a continuous and steady growth trend. The subsystem “green growth” index increased from 0.23 in 2010 to 2.28 in 2019, with a significant increase. The development index of “green welfare” subsystem increased from 0.28 in 2010 to 2.61 in 2019, showing an outstanding growth, which reflecting the improvement of economic development, local public services and residents’ social welfare have also been improved. From this indicated that during the research period, intensive production, recycle of local level improved, however, because the contradiction between the increasing load of resources and environment and the demand of continuously growing economic development exists for a long time, the development of the “green wealth” subsystem, which represents the economy is relatively slow, and it also reflects the difficulty and long-term nature of the sustainable development task of the regional ecological environment system.

The local government make great efforts to promote urbanization, rural revitalization of demonstration area, border central cities of western Yunnan, to promote the production structure, consumption structure, and optimize the elements of sustainable development integration, strengthen local environmental pollution control, reduce the resource consumption gradually, improve the level of regional ecological civilization construction. In addition, from the above model econometric analysis results, it showed that the current level of green development in the region needs to be further improved, and the concept of sustainable development needs to continue to be internalized and integrated into local government decision-making and development planning and design.

#### 3.2. Prediction and calibration of green development index in construction area

The model GM (1, 1) describes the prediction model of the variation trend correlation degree of regional green development index in the short term (Liu sifeng et al., 2000). GM (1,1) contains only one parameter, so it has a large elasticity and wide applicability to measure the sample number and sequence length, and has a good prediction accuracy for local green development system. The multi-index complex system of green development as a single index comprehensive system evaluation by using PPM method, which is substituted into GM (1,1) for model prediction, and the green development trend of the research area in the short term is measured (table 3 below).

**Table3.** Precision grade of Grey Model GM (1,1)

| GM Accuracy-Grade | Small Error-Probability | Posterior Ratio |
| --- | --- | --- |
| Level 1-good | $P \geq 0.95$ | $C \leq 0.35$ |
| Level 2- qualified | $0.80 \leq p < 0.95$ | $0.35 < C \leq 0.50$ |
| Level 3 - fair | $0.70 \leq p < 0.80$ | $0.50 < C \leq 0.65$ |
| Level 4- unqualified | $p < 0.70$ | $C > 0.65$ |

Taking the area as the research object, the characteristic value of green development index (2010-2018) obtained by the above PPM model evaluation and analysis was substituted into the gray model GM (1,1) to predict and test the change of regional green development level in the short term.

Substitute the A= [1.307464273 1.307464209 1.305316214 1.309872736 1.727271031 1.899596651 1.962202329 2.708768395 3.055856406] in GM(1,1), as the initial series x (0), we can get the response function:

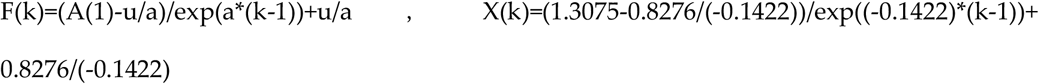

That is X(k)=7.1275/ exp ((−0.1422) *(k-1))-5.82

Thus, the simulation data sequence is generated, and the prediction sequence is obtained through reduction calculation, as shown in Table 4. below.

**Table4.**
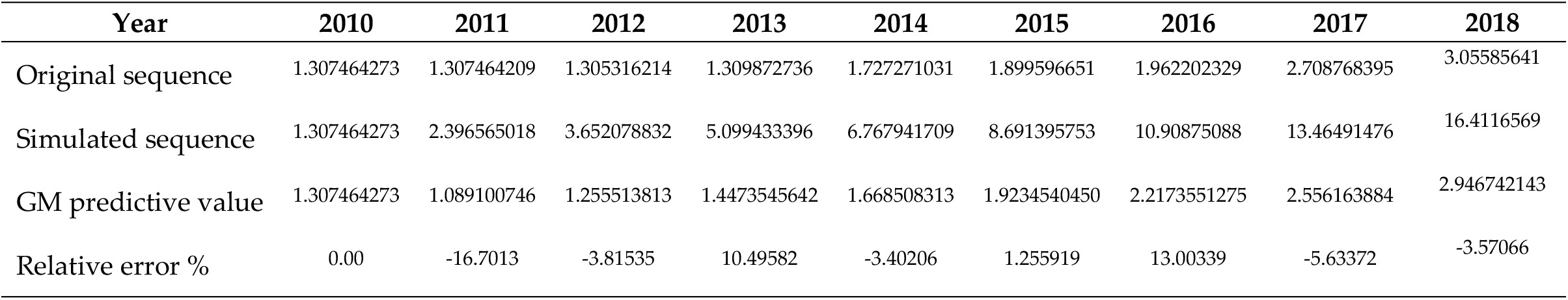
Dynamic evaluation and forecast results of Green Development of Baoshan city Yunnan Province from 2010 to 2018

According to the above, this paper uses the projected eigenvalue 3.708424 of the green development index in 2019 as the verification group, and substitute it into the prediction model to calculate the simulated predicted value of 3.398266, and the relative error is −8.364%. The small error probability P =100%>95%, the posterior ratio C=0.0501<0.35, the average absolute value of relative error −0.9298%<10%, R2 is 0.95, and the prediction accuracy of the model is level 1. Therefore, from 2020 to 2021, the characteristic value of green development in Baoshan 10000 mu project area in Yunnan province can be predicted to be 3.916436 and 4.514896, with an estimated annual growth rate of 15.1%.

### Analysis on Influencing Factors of Regional Green Development

This paper analyzes the influencing factors between the regional green development system and its constituent subsystem index, combined with the situation and problems faced by the comprehensive sustainable development in Baoshan, Yunan Province, and analyzes the interaction and evolution between the green development composite system and subsystems, which are mainly affected by various environmental conditions, such as internal and external conditions. The measurement results show that the local green development index and its composition are mainly affected by environmental pollution, industrial structure, urbanization, population, market and other factors from the three subsystems of ecology, economy and social welfare.

#### (1) Regional Eco-environmental Factors and Green Development

In the process of using PPM model to process the “green wealth” parameters representing the regional ecosystem subsystem, the maximum projection value is 0.455, thus, it can be concluded that the closer to the maximum projection value, that is, the parameters with relatively greater impact on the green development of green wealth subsystem are in the following order: “NPK and compound fertilizer (chemical fertilizer) application amount *(0.56)”, annual afforestation area (0.53), industrial wastewater emission* (0.47), SO2 emission *(0.42), among which “annual afforestation area” is positively correlated with the development and evolution of green wealth subsystem, while other parameters are negatively correlated with it. The green wealth subsystem dropped from the peak value of 1.81 in 2010 to the minimum value of 0.29 in 2016, with a significant decline in development trend, and a slow recovery trend since 2017. With the continuous promotion of local urbanization, the demand for green development and concept of ecological civilization are gradually strengthened; People’s demands for a good ecological environment also promote the local government to improve the ecological environment governance system. For example, the annual afforestation area has increased year by year, and the environmental pollution sources have been well controlled; On the other hand, under the condition of sufficient supply of environmental financial funds, even during 2013-2017, a large area of demolition work was carried out in Baoshan from 2013 to 2017, which inevitably produced a lot of vacant land, wasteland and barren hills. In 2016, the government implemented a construction project of 10000 mu, and the ecological restoration affect gradually showed. As a result, the green wealth sub-index gradually rebounded to 1.1 in 2019, as shown in Figure 3.

**Figure3.**
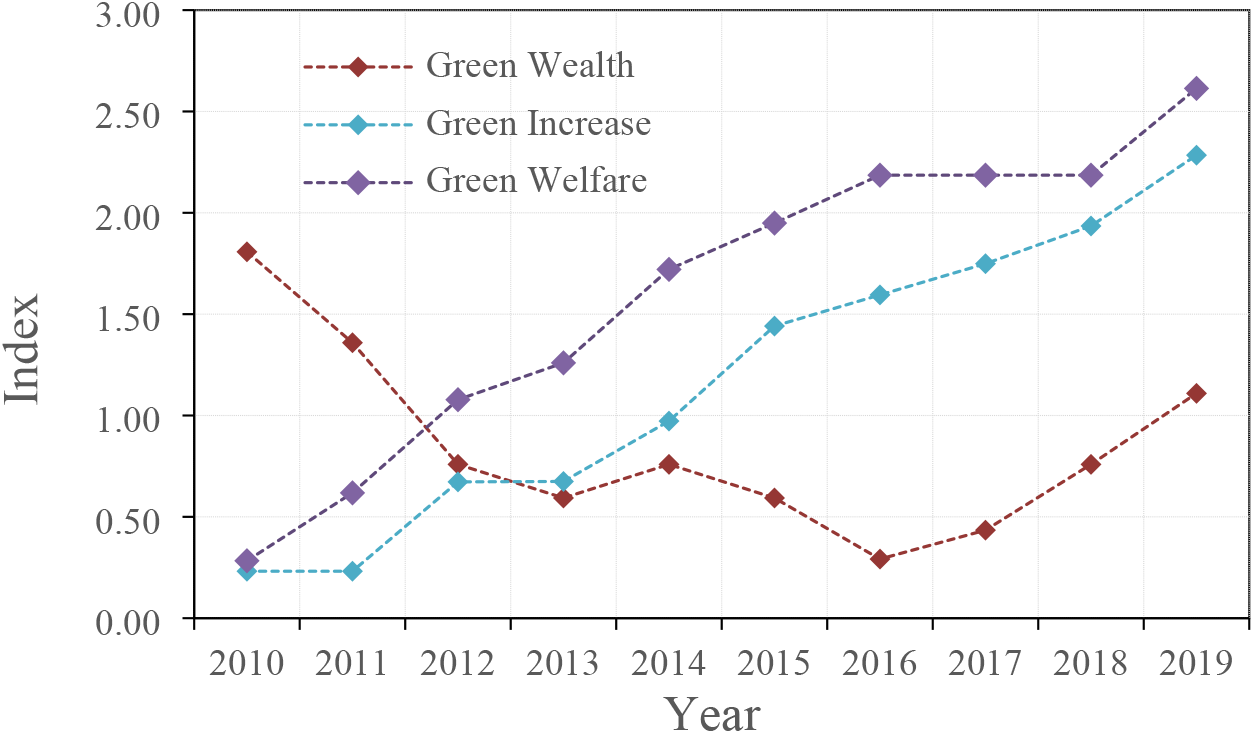
Subsystem index of Green-development of Baoshan’s ecological engineering Area

#### (2) Regional Economic Factors and Green Development

The “green growth” subsystem and its indicators are used to characterize the elements of regional economic development. From Figure 2, the green growth index showed a fluctuating upward trend from 2010 to 2019. According to the research and measurement, the maximum projection value of the green growth subsystem is 0.73, and the parameters that have a great impact on the development of the subsystem include: “proportion of resident energy consumption (0.67)” “energy consumption (t standard coal) (0.45)” “annual output value of the tertiary industry (0.35)” “annual output value of the secondary industry (0.29)” “land output rate (0.28)” “output value of the primary industry (0.24)". Energy consumption is negatively correlated with local economic development, and the others is positively correlated. With China’s rapid economic growth and urbanization process, the total domestic energy consumption of Chinese residents shows an upward trend, accounting for 13% of the total energy consumption, which is lower than the world average level of 31% (Yang Jie et al., 2020), while the maximum proportion of residents’ energy consumption in this study area is 20.3%, which shows a gradual upward trend. This reveals that energy conservation of residents in this region should be one of the key work of energy conservation during the 14th Five Year Plan period, and local governments need to advocate residents’ energy conservation, green consumption and low-carbon life.

In view of the positive related parameters affecting regional green growth, on the one hand, under the background of the current shortage of land resources, making rational use of land, improving land use efficiency and realizing the sustainable development of regional economy, society and ecology turns an important issue studied by local government. The rate of land output is a comprehensive economic index indicating the level of agricultural productivity in a region. On the other hand, urbanization development produced agglomeration effect and spatial spillover effect on local production factors, resulting in synergy and integration between local industries and enterprises, improved input-output efficiency, and gradual optimization and upgrading of industrial structure. With the continuous integration and development of local primary, secondary and tertiary industries, improve the comprehensive development quality and synergy of regional land and resources. Also, the industrial activity is the main behavior of regional system and human relations, and all kinds of pollutants control center, the important carrier of the ecological environment and economy. The combination of resources and environment and economy largely determines the degree and type in the region of resource utilization and economic benefits, and the degree of load on the ecological environment system. Under the background of green development and ecological civilization construction, the adjustment and optimal allocation of local industrial structure are the key points to break the resource and environmental constraints and realize the coordination between human and land.

#### (3) Elements of Local Social Development and Green Development

In the process of measuring the green development coefficient of ecological project construction area in Baoshan, the “green welfare” parameter system is adopted to measure the interaction and influence between local social system element and green development. The overall development of the green welfare subsystem shows a fluctuating rising state during 2010-2019.The optimal projection value is 0.959, with the highest projection value among the three subsystems, and also the subsystem with the most abundant representation of the green development index. The main influencing parameters are as follows: “Urban population (0.427) annual per capita disposable income of urban residents(0.424)”, “per capita occupied public green space area (0.406)”, “per capita net income of farmers(0.331)”, “number of population (0.32)”, “effective irrigated area (0.319)” “rural electricity consumption * (0.284)”, “urban green land rate (0.258)”,etc. except for “rural electricity consumption”, other parameters are positively correlated with the development of green welfare subsystem. On the one hand, urbanization increases the degree of population aggregation, provide more sufficient labor force in primary, secondary and tertiary industries, the income of urban residents, and people’s demand and desire for eco-environmental quality will force the government to pay more attention to local resource and environmental problems, improve and enhance the environment quality, attract high-quality human resources, and enhance the residents’ awareness of ecological civilization, which promote the formation of economic development model of local endogenous growth. On the other hand, the larger population density in the region will cause pressure on the utilization of public resources and environmental quality. For example, the increase of electricity consumption and food demand will reduce the per capita resource ownership in the region, reduce the ecological space, and aggravate the situation of regional resources and environment.

Combine ecological project construction of ten thousand mu and improve regional human settlement environment quality. The ecological sensitive areas within 123.3 square kilometers of Baoshan were strictly protected and restored, with 247,500 square meters of new green areas, 37.23 percent of the built-up areas covered by green areas, and 98.4 percent of the central urban area enjoying excellent air quality throughout the year. A total of 460 million yuan was invested to repair 92,000 square meters of damaged roads and sidewalks, and dredge 36 black and smelly rivers. The living environment in urban and rural areas has been greatly improved.

### Conclusion

Under the theoretical framework of regional green development, this paper studies the green development evolution, trend and influencing factors of 10.000 mu ecological engineering construction area in Baoshan, Yunnan Province by using econometric analysis methods such as projection pursuit evaluation model(PPM) and grey system prediction model, and obtains the following conclusions:

1. The theoretical framework of regional green development is based on the interaction and coordination mechanism of ecological, economic and social complex system of sustainable development from the perspective of system science. Green development is a process of promoting the harmonious development between human and land, human and nature, and the continuous optimization of its development model. Green development covers three subsystems: green wealth, green growth and green welfare. Green wealth is the foundation of local green development construction, green growth is the core of regional economic development, and green welfare is the goal of regional green development system. The coordinated and balanced development of the three subsystem is the internal requirement of green development.
2. As for the temporal evolution of green development, firstly, the overall system index of green development in Baoshan Showed an upward trend, rising from 1.31 in 2010 to 3.71 in 2019, with an average annual growth rate of 18.3%. The temporal evolution of each subsystem shows that the development trend of green wealth and green development and other subsystem is dislocated, indicating that the economic and social development of the study area strongly linked to the development and utilization of ecological resources and environment.
3. The factors that affect the level of regional green development are characterized by locality, dynamics and complexity. The situation of local urbanization, industrial structure, population, environment and resources influence regional green development through economic ecology, land resources development, input-output mechanism.

Based on the requirements of green development in China and the actual situation of regional ecological-economic-social development, the study puts forward the following countermeasures and suggestions:

1. Firmly establish the concept of Chinese ecological civilization and green development, and realize the comprehensive decision-making of local sustainable development. The concept of “GDP only” should be gradually changed, and the concept of coordinated development of man-land system should be internalized into the development planning and decision-making of local governments, the production practice of enterprises, and the whole process of life and consumption of the public. Further strengthen green and economical lifestyles and incorporate ecological civilization education into the teaching content of local colleges, middle schools and primary schools.
2. Improve the level of local ecological economic development and improve the quality of economic development. Take economic ecology as the goal, explore the green, low-carbon, circular and clean development path of traditional industrial economy, promote the formation of circular development system of enterprises, parks, industries and regions, and gradually form a circular development system and integrated management mechanism. Take agricultural supply-side structural reform as the core, ecological engineering construction projects as the main carrier, innovate the main business model. Based on the industry of “vegetable, flower and fruit”, the company innovates three business cards of “Flower Cloud, Fruit Cloud and Vegetable Cloud”, 13,000 mu of vegetables, flowers and fruits have been planted. Meanwhile jointly create smart agricultural science and technology demonstration zone including “modern high value-added agriculture”, “the future of agriculture and incubator”, “balanced nutrition and health experience”, “big data, Internet and electronic commerce” and “public welfare and education”, five new modules, which realize great achievement for the development of modern new agriculture. Accelerate the transfer of surplus land for 10 thousand mu of agricultural ecological tourism zones; play the role of modern enterprise such as East Garden, Huada Genetics Group and Baonong Science and Technology; to develop new types of economic entities, such as professional cooperatives, individual contractors, and mixed joint-stock investment enterprises; to develop large-scale “order” agriculture, provide farmers with all-round technical training, and make modern agriculture more standardized, refined and efficient.
3. Improve the environmental management system, improve the carrying capacity of regional ecological environment, improve the quality of living environment and happiness. Implement the control of regional environmental ecological risk process, implement the protection and restoration supervision of mountains, water, forests, grass, fields and lakes, and improve the zoning scheme of ecological functions. To formulate local development plans for resource conservation and recycling, promote green energy conservation and gradient utilization of energy in local buildings and transportation; Guided by the market construction of resource and environment property rights, ecological compensation system and environmental economic system should be established to reflect the cost of resource and environment depletion and ecological restoration. Supervise the disordered expansion of development areas, gradually optimize the “production-life-ecology space”, and improve the benefits of regional sustainable development; build a system framework that integrates regional urbanization, population, industrial structure, market, resources and environment and other factors to drive the dynamic evolution of green development.

## Author Contributions

For this research articles with 5 authors, “Conceptualization, Qingkui Lai; methodology and analysis, Yunfang Chen and Xuan Ji; writing—original draft preparation, Yunfang Chen; investigation and writing review and editing, Cunyu zhang; data curation, Mengxiong Xiao; project administration, Yunfang Chen. All authors have read and agreed to the published version of the manuscript.

